# Biological Plasticity Rescues Target Activity in CRISPR Knockouts

**DOI:** 10.1101/716019

**Authors:** Arne H. Smits, Frederik Ziebell, Gerard Joberty, Nico Zinn, William F. Mueller, Sandra Clauder-Münster, Dirk Eberhard, Maria Fälth Savitski, Paola Grandi, Petra Jakob, Anne-Marie Michon, Hanice Sun, Karen Tessmer, Tilmann Bürckstümmer, Marcus Bantscheff, Lars M. Steinmetz, Gerard Drewes, Wolfgang Huber

## Abstract

Gene knockouts (KOs) are efficiently engineered through CRISPR-Cas9-induced frameshift mutations. While DNA editing efficiency is readily verified by DNA sequencing, a systematic understanding of the efficiency of protein elimination has been lacking. Here, we devised an experimental strategy combining RNA-seq and triple-stage mass spectrometry (SPS-MS3) to characterize 193 genetically verified deletions targeting 136 distinct genes generated by CRISPR-induced frameshifts in HAP1 cells. We observed residual protein expression for about one third of the quantified targets, at variable levels from low to original, and identified two causal mechanisms, translation reinitiation leading to N-terminally truncated target proteins, or skipping of the edited exon leading to protein isoforms with internal sequence deletions. Detailed analysis of three truncated targets, BRD4, DNMT1 and NGLY1, revealed partial preservation of protein function. Our results imply that systematic characterization of residual protein expression or function in CRISPR-Cas9 generated KO lines is necessary for phenotype interpretation.

## Introduction

Genetic engineering is invaluable for biological research and holds great promise as therapy of genetic diseases. The discovery and development of the clustered, regularly interspaced, short palindromic repeat (CRISPR) technology^1–3^ has made editing widely accessible. This technology is based on an RNA-guided nuclease, such as Cas9 from *Streptococcus pyogenes* bound by a single guide RNA (sgRNA) that provides sequence specificity^4^.

One of the most commonly used CRISPR-Cas9 applications is the generation of a genetic knockout (KO) by the introduction of a frameshift mutation. In this application, the Cas9-sgRNA complex is targeted to a specific sequence in a gene’s coding region and cleaves both strands of the DNA^5,6^. The DNA double-stranded break is repaired by non-homologous end joining, an error-prone pathway introducing insertion or deletion mutations that can lead to frameshifts^7^. A frameshift causes a premature termination codon (PTC) in the expressed transcript, leading to nonsense-mediated decay (NMD) of the mRNA and aberrant peptide products that are degraded^8^.

Multiple studies investigated potential off-target effects of CRISPR-Cas9^9–13^. Collectively, these studies report sgRNA-dependent off-target effects at rates that can be comparable to on-target editing. These findings triggered developments to improve CRISPR-Cas9 specificity by the use of Cas9 nickase-variants^12,14,15^, improvements in sgRNA design^12,13,16,17^, and protein engineering of the nuclease^18,19^.

The editing efficiency of CRISPR-Cas9 is commonly assessed on the genomic level. However, many current study designs do not further assess whether the induced frameshift mutation results in the expected complete loss of protein expression and activity. Here, we systematically characterized the effects of frameshift KO mutations. We studied 193 isogenic KOs in HAP1 cells by quantitative transcriptomics and proteomics. Although all KOs contained frameshifts, about one third of them still expressed the target protein, albeit the majority at reduced levels. Similarly, we detected residual target expression for a CRISPR-Cas9 KO in a K562 cell line. A detailed analysis of three of the KO lines (BRD4, DNMT1 and NGLY1) revealed that the detected protein products were truncated but preserved partial functionality.

## Results

### Systematic characterization of frameshift KO mutations

We investigated 193 HAP1 lines, in each of which one of a set of 136 genes was targeted with a frameshift KO mutation by CRISPR-Cas9. All lines were derived from the near-haploid human cell line HAP1, which contains a single copy of most genomic loci and is an ideal model system to trace genome-editing events^20–22^. In a first set of experiments, RNA and protein expression levels of 174 of the mutants and the parental line were measured using 3’RNA-sequencing^23^ (3’RNA-seq) and triple-stage mass spectrometry (MS3) with a single replicate for each line (**Fig. 1a**). First, we looked in each KO line at the transcripts originating from the targeted gene and calculated the ratio between the expression level in the KO and the parental line, which we termed *residual level*. A residual level of 0 indicates complete elimination, a value of 1 indicates a target transcript level as high as in the parental line. Contrary to our expectation, the 174 residuals levels did not concentrate near 0, but covered a wide range. Thus, in many KO lines the mutant mRNA was not strongly reduced. Since in all lines the presence of a premature termination codon was confirmed by DNA sequencing, this implies that there was no or weak NMD response to these transcripts. This finding is in line with a recent study that reported high variation in NMD efficiency depending on a complex set of biological factors^24^.

**Figure 1:**
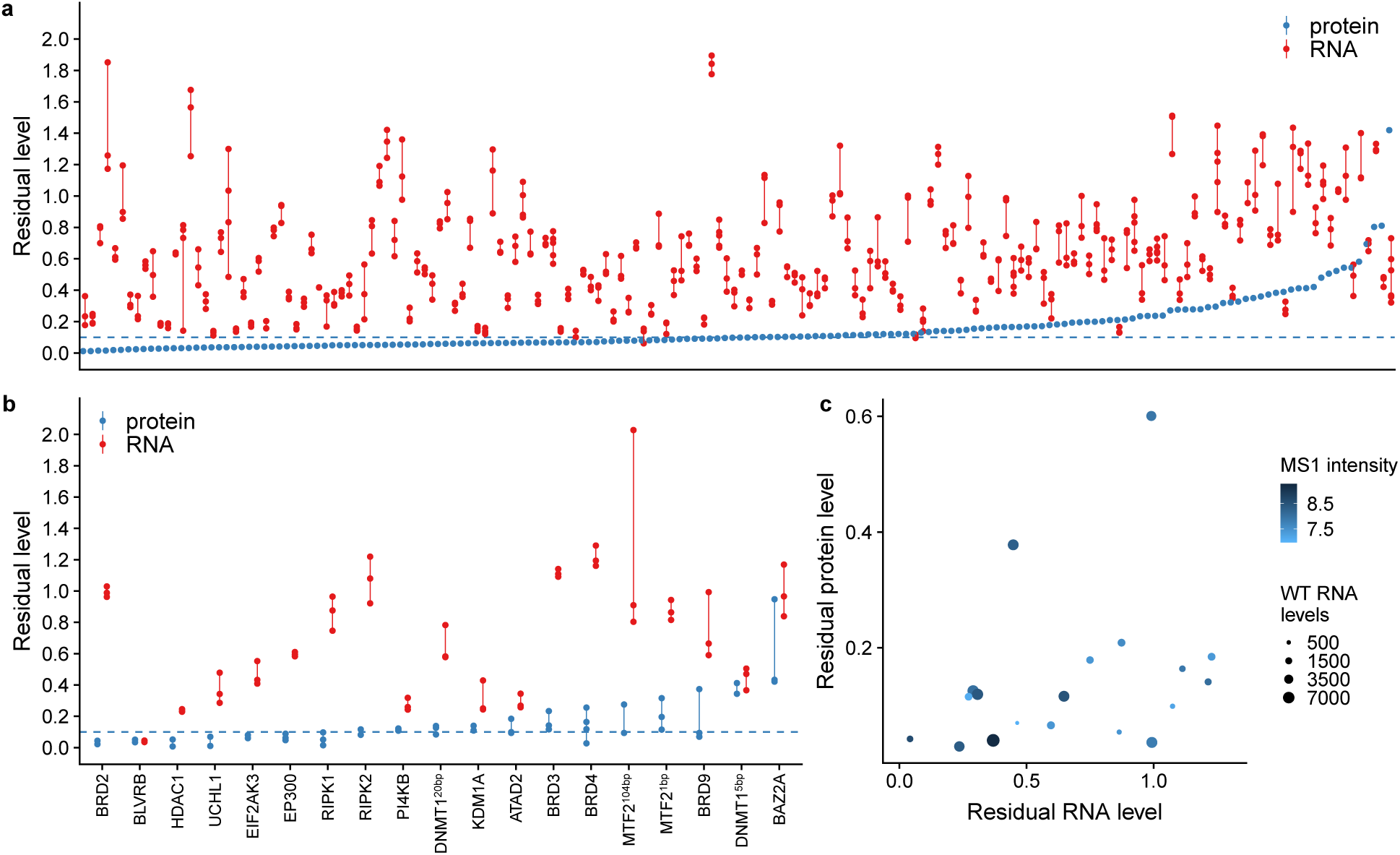
Residual transcript and protein expression in a set of 193 HAP1 cell lines harbouring frameshift KO mutations. (**a**) Residual RNA and protein levels for 174 of the lines. RNA levels were quantified with 3’RNA-sequencing with two or more biological replicates, indicated by red points connected by vertical lines. Protein levels were measured with a single MS3 replicate (blue points). (**b**) Residual RNA and protein levels of the remaining 19 lines, which we quantified by whole-transcript RNA-sequencing and MS3, each with two or more replicates. The data are presented as in Panel a. (**c**) Scatterplot of RNA and protein expression levels for the 19 KO lines displayed in Panel b. Size of dots indicates the WT RNA level. Color indicates the protein MS1 signal intensity.

Next, we performed an analogous analysis for target protein levels. To this end, we first benchmarked our method, tandem mass tags (TMT) isobaric labeling in combination with MS3 analysis^25^, with a control experiment in which we spiked *E. Coli* into human protein background in different ratios (**Fig. S1a and S1b**). Consistent with previous results^25^, quantification of lowly expressed *E. Coli* proteins (≤ 20% of the human level) was more accurate with the MS3 method compared to standard tandem mass spectrometry (MS2) (**Fig. S1c**). To assess specificity, we considered the distribution of *E. Coli* protein quantifications in a human-only sample. While ideally these should all be zero, for 8% (49 out of 587), the quantification exceeded 0.1 of the value in the *E. Coli*-only sample (all of these were based ≤ 8 unique spectrum matches; **Fig. S1f and Table S2**). Based on this, we chose a quantification ratio between KO and parental line of 0.1 as a threshold for calling residual expression of the targeted protein and assumed that this would imply a false positive rate around 8% (**Fig. S1d and S1e**). For all targeted proteins, the quantification was robust, with at least 2 unique peptides in at least one of the 10 multiplexed samples of the corresponding TMT experiment. Surprisingly, about one third of the KO lines displayed residual protein levels around 10% or higher (**Fig. 1a**). Moreover, there was no clear correlation between residual RNA and protein levels, indicating that residual RNA levels are not predictive of residual protein levels.

We selected an additional set of 19 lines for quantification with whole transcript RNA-seq and MS3 proteomics (**Fig. 1b**). Each of these quantifications was done with two or more replicates, and replicates were highly reproducible (**Fig. S2 and S3**). Similar to the observation in Figure 1a, there was little correlation between residual RNA and protein levels. Moreover, residual RNA and protein levels were not dependent on the initial levels prior to CRISPR modification, as indicated by number of reads and MS1 intensities (determined by the TOP3 method^26^) of the parental samples (**Fig. 1c**).

### Residual protein expression due to exon skipping

The MTF2^104bp^, PI4KB and BRD3 KO lines showed residual protein levels ranging from 9% to 28% (**Fig. 1b**). The transcript level of PI4KB was reduced, whereas MTF2^104bp^ and BRD3 displayed levels close to WT. Hence we assessed if the mutant lines express alternative transcript forms which skip the edited exon (**Fig. 2**).

**Figure 2:**
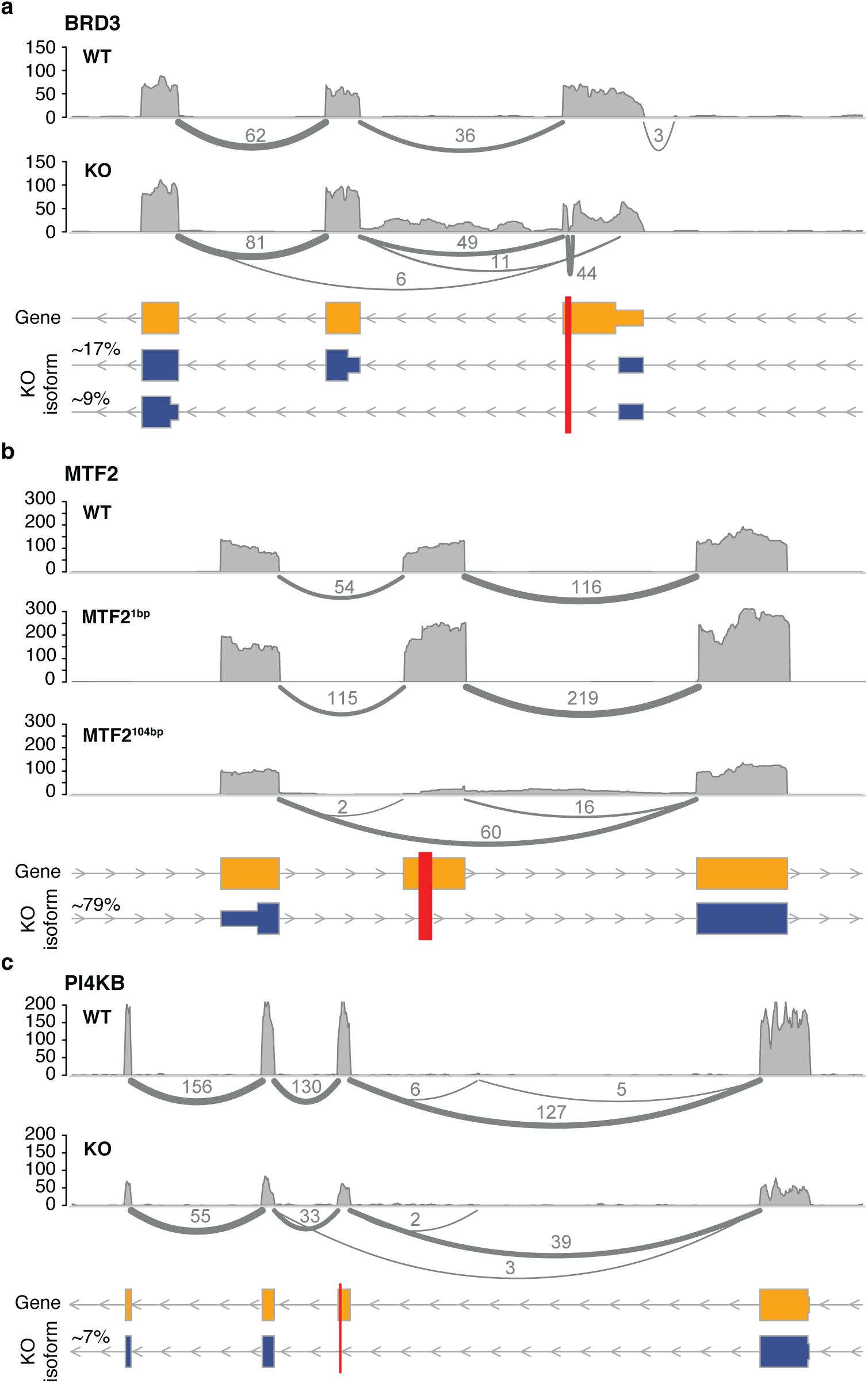
Residual protein expression due to exon skipping. Sashimi plots indicate exon usage and splicing of the BRD3 (**a**), MTF2 (**b**), and PI4KB (**c**) transcripts in their respective KO lines. Arches and numbers represent exon-exon junction RNA-seq reads. The canonical isoforms are shown in yellow, alternative isoforms identified in the KO lines in blue. Red bars indicate the sgRNA target sites.

The *BRD3* transcript in the BRD3 KO line contained exon-exon junction reads spanning from within the 5’ untranslated region (UTR) to the second coding exon (11 out of 66 reads mapping to this gene) and from within the 5’ UTR to the third coding exon (6 out of 66 reads) (**Fig. 2a**). This splicing event removes both the start codon and the frameshift mutation from the transcript. Our data suggest that the 5’ UTR is extended and Methionine (Met) 81 or Met 125 are used as alternative start codons giving rise to a BRD3 protein that is N-terminally truncated.

In the MTF2^104bp^ line, the 104 bp insertion and associated frameshift induced a strong alternative splicing event (**Fig. 2b**). Exon 5, which contains the frameshift mutation, is skipped within the transcript (60 out of 76 reads). Strikingly, this did not occur in another line generated using the same sgRNA, where MTF2 was mutated with a 1 bp insertion (**Fig. 2b**). Deletion of exon 5, which is 101 nucleotides long, will lead to a frameshift downstream of the exon. Interestingly, there is an out-of-frame ATG in exon 4 (Chromosome 1: 93,114,751-93,114,753), which after skipping of exon 5 would result in an in-frame transcript from exon 6 on. This could explain the observed residual protein expression in the MTF2^104bp^ line, with a residual protein isoform that omits the original first 160 amino acids and instead starts with 10 alternative amino acids originating from the latter part of exon 4

The PI4KB frameshift mutation resulted in low level expression of a splice variant skipping the frameshift-containing exon 5 (**Fig. 2c**), causing an in-frame deletion of 228 bp (76 amino acids). Estimated from the number of exon-exon junction reads, this splicing event only occurred in ∼7% of PI4KB transcripts (3 out of 42 reads). This suggests that the low-level residual protein originates from the alternatively spliced transcript.

### BRD4 frameshift mutation causes a partially functional BRD4 truncation by translation reinitiation

The BRD4 KO line contained a 16 bp deletion in exon 2 (**Table S1**) and showed residual RNA and protein expression (**Fig. 1b**). However, we did not find evidence for exon-skipping (**Fig. 3a**). To further characterize this mutant, we studied the residual protein in more detail. We used the data from the quantified tryptic peptides to get an overview of residual expression of different regions of BRD4. The mutant expressed low levels of the N-terminal protein region, corresponding to the exons before and around the CRISPR-Cas9 target site (vertical dashed line) (**Fig. 3b**). The protein part C-terminal of the target site displayed residual expression levels higher than 10%. These data suggest N-terminal truncation of BRD4 as a consequence of translation reinitiation^24,27,28^. We confirmed the residual expression of BRD4 by Western blotting (**Fig. 3c**). BRD4 appeared to be absent from the BRD4 KO line at standard exposure levels, but extended exposure revealed a smaller protein product with an intensity of approximately 15%. To further validate these results, we performed BRD4 immuno-precipitations (IP) with two different BRD4 antibodies (**Fig. 3c**). Both the WT and the truncated product were enriched by both antibodies, confirming the BRD4 identity of the truncated protein.

**Figure 3:**
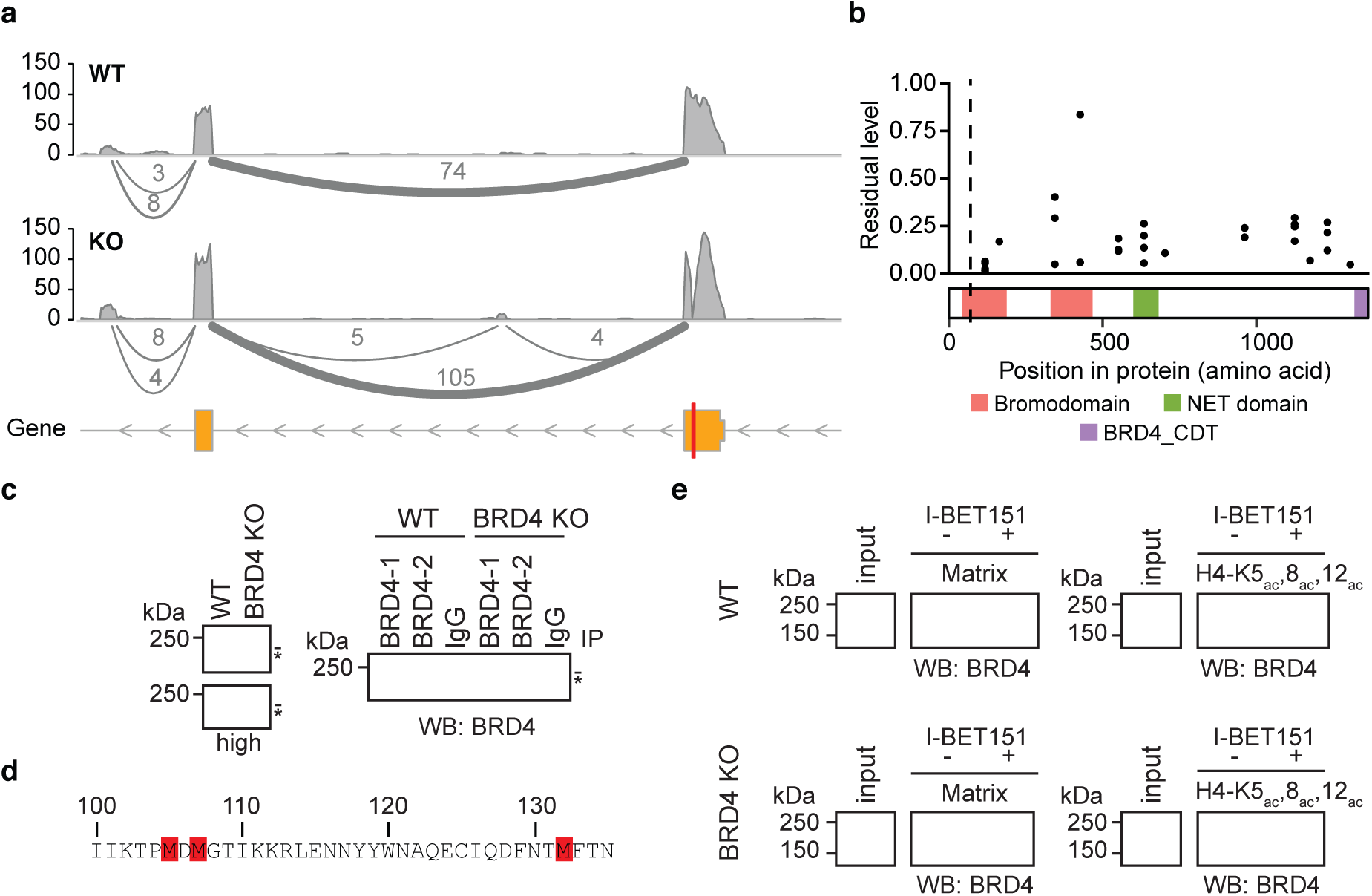
Residual expression of a partially functional, truncated BRD4 protein due to translation reinitiation. (**a**) Sashimi plot indicating exon usage and splicing of BRD4 in the BRD4 KO line. Layout and colors as in Figure 2. (**b**) Residual peptide levels, aggregated per exon. For each biological replicate, the median of all spectra assigned to the amino acid sequence of each exon is depicted. The dashed vertical line depicts the sgRNA target site. The protein’s domain structure is shown at the bottom of the plot. (**c**) Assessment of the BRD4 truncation by Western blotting (WB). Left panel: BRD4 staining of WT and BRD4 KO inputs with normal (top) and high (bottom) exposure. Right panel: BRD4 IPs with two BRD4 antibodies (BRD4-1 and BRD4-2). (**d**) Amino acid sequence of position 100-135 of BRD4, highlighting the methionines that can act as translation reinitiation sites. (**e**) Binding studies with cell extracts expressing WT or truncated BRD4. Affinity pulldowns using beads with immobilized inhibitor (panBET matrix) (left panels) or biotinylated histone H4ac tails (right panels) for the parental (top panels) and BRD4 KO (bottom panels) line. Matrix: Beads with immobilized panBET. I-BET151: bromodomain inhibitor. H4-K5_ac_,8_ac_,12_ac_: acetylated histone H4 tail at positions K5, 8 and 12.

The abundance levels of the quantified peptides and the size of the truncated protein support the explanation that translation of the protein is reinitiated at Met 105, 107 or 132 (**Fig. 3d**). Notably, Met 105 is highly conserved across the bromodomain protein family^29^. For BRD2, a putative isoform is described starting at the methionine corresponding to Met 105 in BRD4^30^. Additionally, one of the identified BRD3 truncations (**Fig. 3d**) starts at Met 81, which also corresponds to Met 105 in BRD4. Altogether, these properties of Met 105 support its ability of initiating an N-terminal truncated protein product. The truncated isoform lacks the initial 30 amino acids (41%) of the first bromodomain (BD1). BD1 is crucial for binding of BRD4 to acetylated histone H4^29^ and is thought to mediate the cellular function of BRD4 as a transcriptional regulator. To test whether the truncated protein retained any functionality, we performed binding assays with a histone H4 N-terminal peptide and with a small molecule bromodomain inhibitor (I-BET). I-BET binds to the bromodomains of BRD4 and other BET proteins with similar affinity^31,32^. Sepharose beads derivatized with I-BET were capable of binding both WT and truncated BRD4 as expected, since the truncated protein still comprises the second bromodomain (BD2) (**Fig. 3e**, left panels). In contrast, the triple acetylated Histone H4 tail peptide, binding to BD1, showed binding to WT but not truncated BRD4 (**Fig. 3e**, right panels). These results confirm the loss of BD1 and the preservation of a functional BD2 in the BRD4 KO.

### DNMT1 frameshift mutation leads to retained DNMT1 functionality

Two DNA methyltransferase 1 (DNMT1) KO lines were measured with whole-transcript RNA sequencing and MS3-based quantitative mass spectrometry with 2 (DNMT1^5bp^) and 3 (DNMT1^20bp^) replicates (**Fig. 1b**). The DNMT1^5bp^ line has a 5 bp deletion in exon 6 (**Table S1**) and shows residual peptide levels of about 40% nearly uniformly throughout the gene body (**Fig. 4b**). Although we observed three reads supporting skipping of the 79 bp long CRISPR-targeted exon (**Fig. 4a**), this would still lead to a frameshift in the spliced transcript. Instead, the expression pattern at the peptide level is better explained by two truncated proteins. The induced 5 bp deletion results in a stop codon directly downstream of the CRISPR target site, leading to a C-terminally truncated protein product consisting only of the first 188 amino acids. This is validated by the complete absence of the “RKPQEESER” peptide (amino acids 178-186) in the KO but not the WT line. Although the peptide is still present in the truncated protein product, its new location close to the C-terminal end (RKPQEESERAKstop) results in a modified cleavage context for trypsin. Thus, the peptide is no longer preferentially generated and quantified. A second, N-terminally truncated protein is explained by intron retention. The first quantified peptide downstream of the CRISPR target site is located at the end of exon 8 (peptide “EEERDEK”, amino acids 234-240). Retention of the intron between exons 7 and 8 creates three in-frame methionines that could be used for translation reinitiation. This hypothesis of intron retention is in line with existing annotation of DNMT1 with twelve transcripts that contain retained introns^33^.

**Figure 4:**
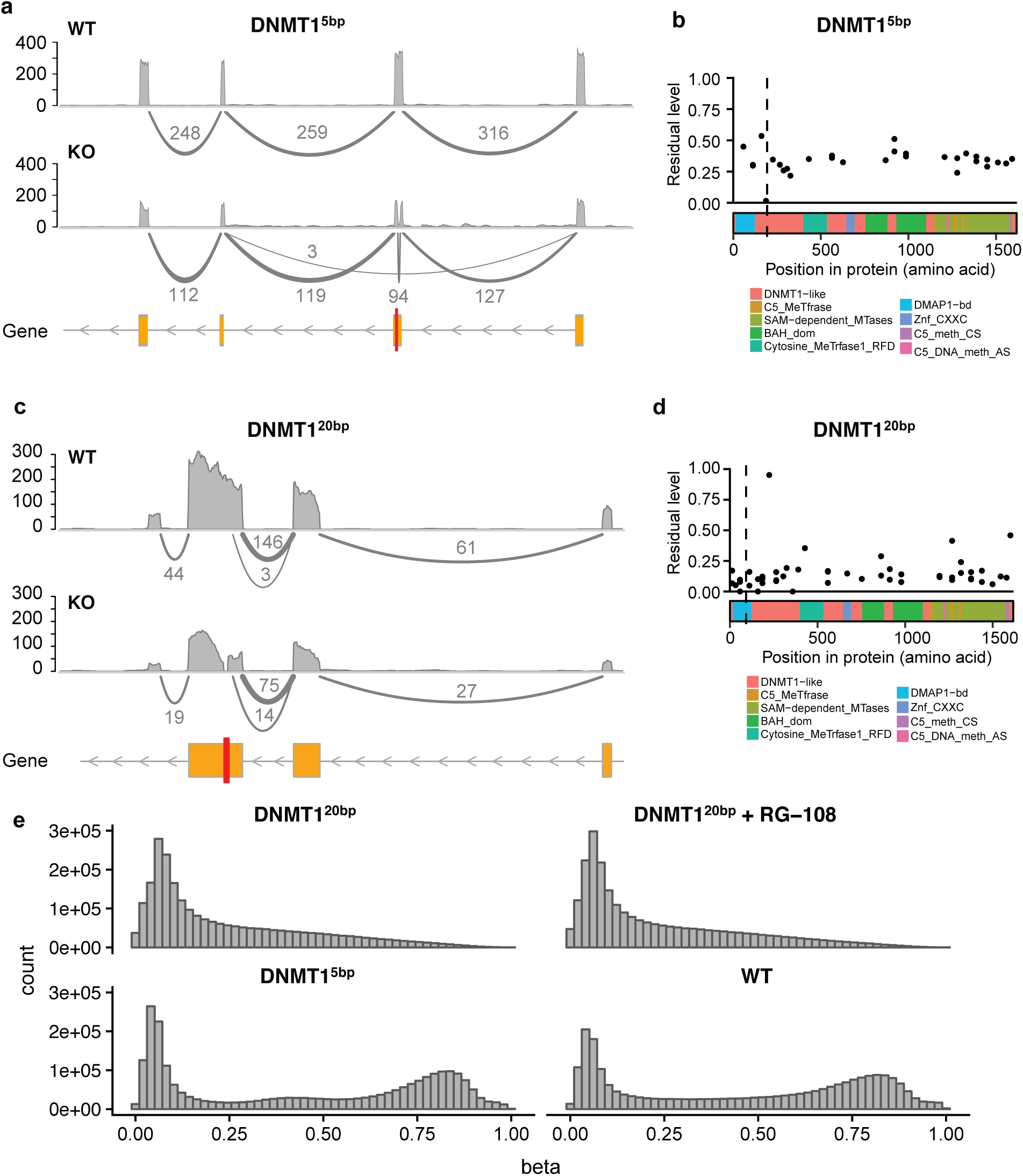
Residual protein expression results in retained activity of DNMT1 in one of the two KO lines. (**a and c**) Sashimi plot indicating exon usage and splicing of DNMT1 in the DNMT1^5bp^ and DNMT1^20bp^ lines, respectively. Layout and colors as in Figure 2. (**b and d**) Residual levels of exons at the protein level of DNMT1^5bp^ and DNMT1^20bp^, respectively. (**e**) Histograms of *β*-values from a methylation array experiment. For each pair of probes binding to a given CpG site, *β* = *M/*(*M* + *U*) where *M* and *U* are the intensities of the methylated and unmethylated probes.

The DNMT1^20bp^ line has a 20 bp deletion in exon 4. For this line, we did not find evidence for excision of the targeted exon (**Fig. 4c**). The nearly uniform residual peptide level of about 10% throughout the gene body (**Fig. 4d**) is at the noise threshold established in the MS3 benchmark (**Fig. S1**).

To assess residual protein activity of both mutant lines, we measured genome-wide methylation levels using methylation arrays. DNMT1, the main protein involved in maintenance of DNA methylation, copies pre-existing methylation marks onto hemimethylated DNA strands during DNA synthesis^34^. The DNMT1^5bp^ line shows methylation levels similar to WT (**Fig. 4e**), consistent with residual protein activity. In contrast, the DNMT1^20bp^ line shows weak to no evidence of DNA methylation (**Fig. 4e**). Moreover, treating the DNMT1^20bp^ line with the DNMT1 inhibitor RG-108 does not change its methylation levels, consistent with already complete abrogation of DNMT1 methylation activity. These results show that one of the two investigated DNMT1 KO lines, DNMT1^5bp^, has residual protein activity.

### NGLY1 frameshift mutation causes a partially functional NGLY1 truncation

To extend our observations to a second cell type, we assessed frameshift KO mutations of the *NGLY1* gene, which encodes a de-*N* -glycosylating enzyme, in K562 cells (**Fig. 5**). Two KO clones termed NGLY1^c15^ and NGLY1^c20^, that resulted from expansion of a single cell, were isolated to enable comparison of different frameshift mutations. By MS3-based quantitative mass spectrometry, we did not detect residual target protein expression in the KO lines (**Fig. 5a**). Interestingly, RNA sequencing revealed skipping of exon 3, which in both clones contained the frameshift mutation (**Fig. 5b**). Skipping of this exon results in an in-frame deletion of 246 bp, leading to a truncated protein missing Glycine 74 to Threonine 155. Although not detected by mass spectrometry, the expression of the in-frame exon-3-skipping transcript led us to investigate potential residual function of the target enzyme. To characterize the deglycosylation activity of NGLY1, we performed a deglycosylation-dependent Venus fluorescence assay followed by cell sorting^35^. Both NGLY1 KO lines showed deglycosylation activity, albeit reduced to 60% to 65% compared to WT cells (**Fig. 5c**). Interestingly, the addition of the NGLY1 inhibitor zVAD-FMK^36^ further reduced the Venus fluorescence in the two NGLY1 mutant lines (**Fig. 5c**). To rule out nonspecific effects of zVAD-FMK, we generated patient-derived^37^, termed m01, and frameshift, termed m03 and m14, mutations in NGLY1 exon 8, that completely abrogated deglycosylation activity (**Fig. 5c**). Indeed, zVAD-FMK treatment did not further reduce Venus fluorescence in the exon 8 edited clones (**Fig. 5c**). These results indicate that the cells of the clones NGLY1^c15^ and NGLY1^c20^ had reduced but measurable NGLY1-based deglycosylation activity.

**Figure 5:**
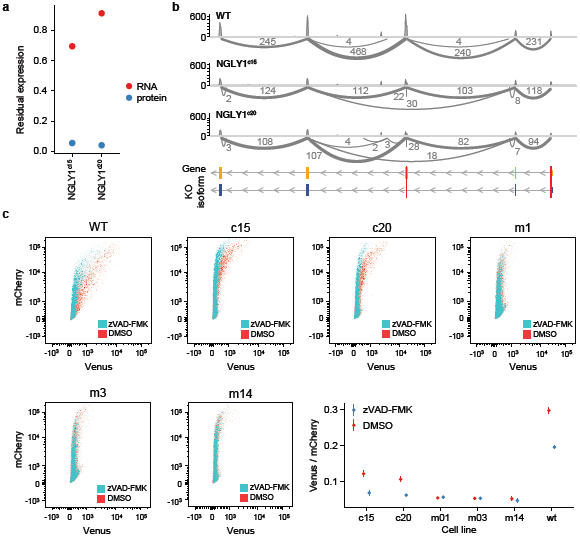
NGLY1 frameshift mutation in K562 cells results in a partially functional NGLY1 truncation. (**a**) Residual RNA and protein levels of two NGLY1 frameshift mutation clones. Residual RNA levels were determined from a single RNA-seq experiment, residual protein levels from all spectra assigned to the protein in a single mass spectrometry measurement. (**b**) Sashimi plots indicating exon usage and splicing of the NGLY1 transcript in these and the parental lines. (**c**) Deglycosylation-dependent Venus fluorescence assay results for the WT and NGLY1 frameshift mutation lines stably expressing the ddVenus reporter. The square panels show FACS plots, quantifications are shown in the lower right. zVAD-FMK: NGLY1 inhibitor. c15 and c20: clones with frameshift mutations in exons 1 and 3. m01: clone with a patient-derived^37^ mutation. m03 and m14: clones with a mutation in exon 8.

## Discussion

We measured residual RNA and protein levels of genes targeted by frameshift mutations introduced by CRISPR-Cas9 and non-homologous end joining in 193 HAP1 cell lines corresponding to the KO of 136 target genes. We studied residual protein function for two of these lines. Moreover, we characterized six K562 cell lines with *NGLY1* frameshift mutations by transcriptomics, proteomics and functional assay. The lines differed widely in their residual protein levels and mechanisms underlying residual expression, which we summarize in a model that delineates the possible effects of frameshift KO mutations (**Fig. 6**).

**Figure 6:**
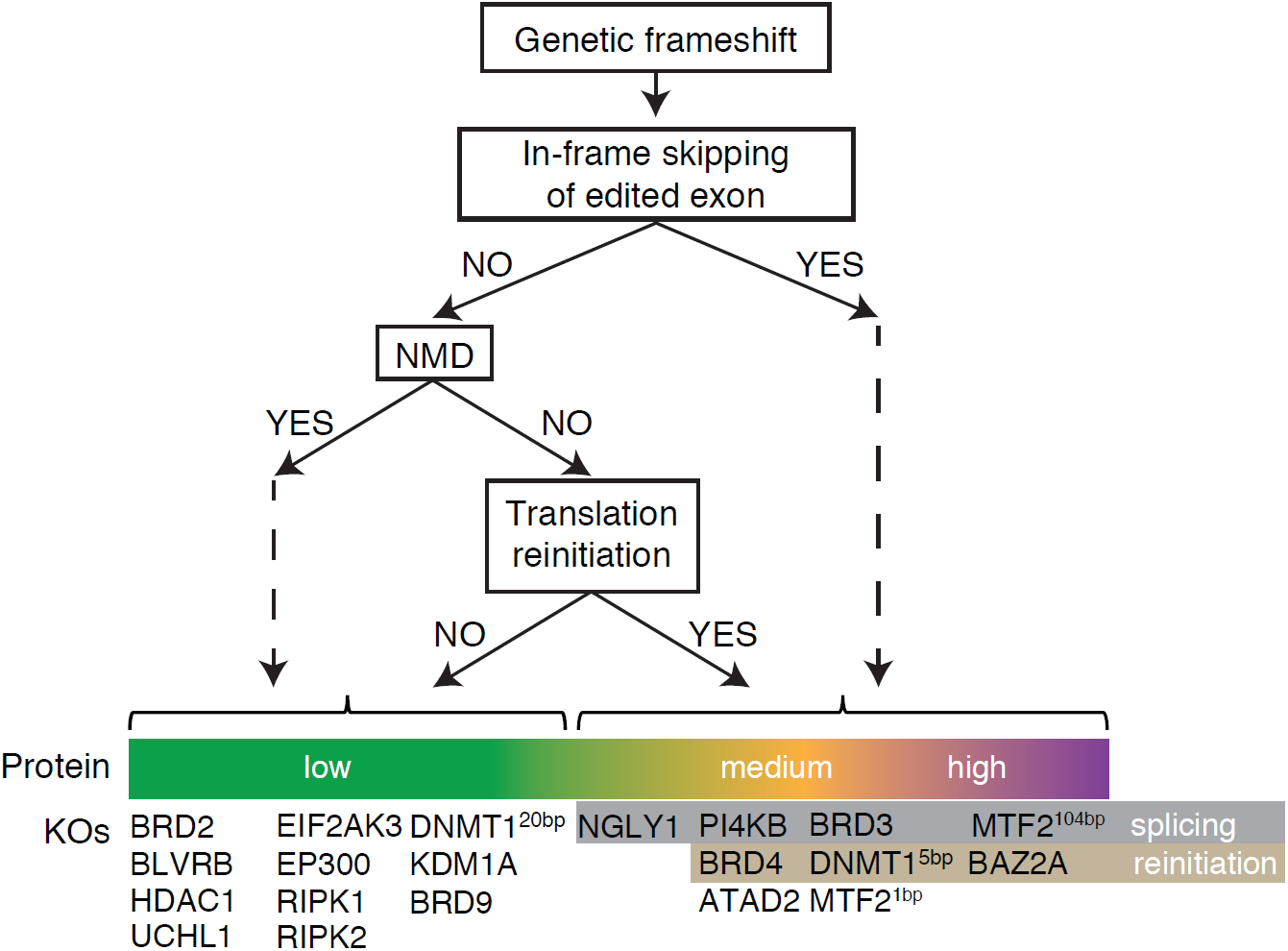
Consequences of CRISPR-Cas9-generated frameshift mutations. Decision tree denoting the possible consequences of a frameshift mutation. At the bottom, resulting protein levels are indicated together with the categorization of the 19 HAP1 KO lines of Figure 1b and the NGLY1 KO in K562 cells.

Our first striking observation was the heterogeneity in NMD efficiency in the KO lines. Both alternative splicing, which deletes the premature termination codon from the transcript, and translation reinitiation appeared as causes for incomplete NMD efficiency. However, even in the lines without residual protein expression, NMD efficiency—as assessed by the residual RNA level—was variable. This observation is in line with a recent study reporting multiple mechanisms influencing NMD efficiency^24^, and indicates that frameshift mutations can lead to full disruption of the protein despite no or weak NMD.

The second striking observation was the residual protein expression in about one third of the KO lines. We identified skipping of the frameshift mutation in the transcript by alternative splicing in four characterized lines. This is in line with previous anecdotal reports^38,39^ and identifies this mechanism as a major source of KO eviction. Moreover, we identified residual protein expression as a consequence of translation reinitiation in three of the KO lines (**Fig. 6**). These findings extend the report of Makino et al^28^ and indicate that translation reinitiation is another unexpected potential effect of CRISPR-Cas9-induced frameshift mutations that has to be controlled for. The production of truncated proteins by alternative splicing and/or translation reinitiation is an important potential limitation of the used CRISPR technology. However, it should be possible to at least partially suppress or circumvent these mechanisms by adapting the experimental design, for instance by using combinations of sgRNAs and/or modifying splicing sites.

We observed that transcript levels of the frame-shifted gene are not predictive of protein levels. This highlights the need for careful and detailed protein-level characterization of frameshift mutations generated by CRISPR technology. The mass spectrometry-based proteome-wide quantification applied here has high signal-to-noise ratio^25^ and can resolve multiple peptides in different regions of a protein. It is more sensitive and less biased than Western blotting, which has lower signal-to-noise ratios and is limited to the epitope of the antibody. As illustrated by the BRD4 KO Western blot (**Fig. 3c**), lower expressed and truncated proteins are easily overlooked. Therefore, we recommend careful characterization of KO lines generated by CRISPR-Cas9 induced frameshifts through high-resolution mass spectrometry.

Residual protein expression of the alternatively spliced NGLY1 frameshift mutation clones was not detected by mass spectrometry, even though residual deglycosylation activity was observed in these lines. This argues for low residual protein expression below the detection threshold of SPS-MS3. We hypothesize that this low amount of enzyme is already sufficient, in the cellular environment, to generate enough enzymatic activity in the time-scale of the applied deglycosylation assay.

At least two of the target genes for which we observed residual protein expression, BRD4 and DNMT1, are known to be essential for viability in HAP1 cells^40,41^. In clones containing frameshift mutations, the essentiality of these genes likely promotes the emergence of splice variants or translational reinitiation events that retain gene function. Although this might undermine the original intentions of the experimenter, especially if the residual protein remains unnoticed, it also enables identification of nonessential domains within essential genes. The remaining functionality of the truncated BRD4 together with the observed unrestricted cell growth indicate that, unexpectedly, the BD1 domain is not essential in BRD4 proteins in HAP1 cells. BD1 is conserved across the BET protein family and mediates chromatin binding. It is tempting to speculate that BD1-dependent functions can be compensated by BRD2 or BRD3, and that the essential function of BRD4 is carried out by the non-conserved C-terminal portion.

In the case of DNMT1, one of our two mutant clones exhibited loss of protein expression to background levels, concomitant with a striking loss of DNA methylation that was not reduced further by an inhibitor drug. Yet this mutant line was viable, which may be explainable by slow emergence of compensatory mechanisms during clone isolation, which were not observed under the conditions of the studies that reported essentiality of DNMT1.

In conclusion, our systematic approach enabled us to study the prevalence and nature of adverse effects of CRISPR-Cas9-induced frameshift mutations on target protein levels. We show that both alternative splicing and translation reinitiation frequently occur in such KO lines, resulting in partially functional protein truncations. Residual protein function of CRISPR-Cas9 frameshift mutations is an important potential limitation of the CRISPR technology for biological research as well as therapeutic applications.

## Methods

### Culturing and sequencing of KO clones

HAP1 KO clones were obtained from Horizon Discovery. Cells were grown in IMDM (GIBCO, #21980-032) supplemented with 10% FCS (GIBCO, #10270) and harvested at 70-90% confluency. Gene target indels from KO cell lines were checked by Sanger sequencing (Sequiserve, Vaterstatten, Germany) from cells with similar passage number as those used to derive transcriptomics and proteomics samples.

### RNA sequencing

For the quantification of RNA levels, two sequencing techniques were employed. In the technique used for the data shown in Figure 1a, each KO line was measured by 3’RNA-sequencing^23^ to a depth of 2 million reads per sample on average. Since this sequencing technique generates reads only at the 3’-end of each transcript molecule, it provides reliable quantification of transcripts at a fraction of the depth that would be needed with shotgun RNA-seq. For the detailed investigation of the subset of 19 KO lines shown in Figure 1b and the analysis of alternative splicing patterns, KO cell lines were measured with whole-transcript RNA-sequencing. This technique generates reads throughout all regions of the gene body, and it resulted in a depth of 27 million reads per sample on average.

#### 3’RNA-sequencing

Total RNA was isolated from 2-3 million HAP1 cells using the Direct-zol-96 RNA kit (Zymo Research) according to the manufacturer’s protocol. RNA concentration and integrity were determined on a Fragment Analyzer (Advanced Analytical). cDNA libraries for 3’ mRNA-seq were prepared from 500 ng total RNA using the QuantSeq 3’ mRNA-Seq Library Prep Kit FWD for Illumina (Lexogen) according to the manufacturer’s protocol. For library amplification 12 PCR cycles were applied and 45-48 libraries were pooled for sequencing on HiSeq 2000 or HiSeq 2500 instruments (Illumina) using 50 base pair single end reads.

#### Whole-transcript RNA-sequencing

Total RNA was isolated from 2-3 million HAP1 cells using the Direct-zol-96 RNA kit (Zymo Research) according to the manufacturer’s protocol. RNA concentration and integrity were determined on a Fragment Analyzer (Advanced Analytical). Preparation of cDNA libraries was done with the TruSeq RNA Library Prep Kit v2 from Illumina according to the manufacturer’s instructions. cDNA was amplified by 15 PCR cycles and 10-11 samples were pooled before sequencing for 100 base pair single-end reads on a HiSeq 4000 instrument (Illumina).

#### Data analysis

RNA-seq reads were aligned with STAR version 2.5.3a^42^ to the Human Reference Genome GRCh37. Mapping of reads to the genome and the presence of frameshift mutations were investigated with IGV^43^. Differential expression analysis was performed using the R/Bioconductor package DESeq2^44^. Sashimi plots were generated using the R/Bioconductor package Gviz^45^ and gene models were obtained from ENSEMBL using biomaRt^46^.

### MS3 proteomics

Cells were harvested by scrapping into cold PBS, washed, and snap-frozen in liquid N_2_. Lysis was done in 2% SDS for 3 min at 95 °C in a thermomixer (Thermo Fisher Scientific), followed by digestion of DNA with Benzonase at 37 °C for 1.5 h. Lysate was cleared by centrifugation and the protein concentration in the supernatant was determined by BCA assay. Proteins were reduced by TCEP and alkylated with chloroacetamide, separated on 4-12% NuPAGE (Invitrogen), and stained with colloidal Coomassie before proceeding to trypsin digestion. Alternatively, proteins were digested following a modified solid-phase-enhanced sample preparation (SP3) protocol^47^. Briefly, proteins in 2% SDS were bound to Paramagnetic beads (SeraMag Speed beads, GE Healthcare, CAT#45152105050250, CAT#651521050502) on filter plates (Multiscreen, Merck-Millipore, CAT#10675743) and filled to 50% Ethanol. Following washing of beads with 4 times 200 *µ*L 70% Ethanol, beads are resuspended in trypsin and LysC in 0.1 mM HEPES (pH8.5) containing TCEP and chloroacetamide and incubated at RT overnight. Peptides are collected and after lyophilisation subjected to TMT labelling.

*E. Coli* spiked into human protein background were produced by a tryptic digest of an *E. Coli* as described above. Peptides were pooled and separated at defined ratios prior to TMT labelling. The human background consisted of pooled tryptic digests from human cell lysates labelled with TMT.

Peptides were labeled with isobaric mass tags (TMT10; Thermo FisherScientific, Waltham, MA) using the 10-plex TMT reagents, enabling relative quantification of 10 conditions in a single experiment^48,49^. The labeling reaction was performed in 40 mM triethylammonium bicarbonate or 100 mM HEPES, pH 8.5 at 22 °C and quenched with glycine. Labeled peptide extracts were combined to a single sample per experiment and lyophilized.

#### Sample preparation for MS, Pre-fractionation

Lyophilized samples were re-suspended in 1.25% ammonia in water. The whole sample was injected onto a pre-column (2.1 mm *×* 10 mm, C18, 3.5 *µ*m [Xbridge (Waters, Milford, MA)]) at a flow rate of 15 *µ*L min^-1^. Separation was done at 40 *µ*L min^-1^ on a reversed-phase-column (1 mm *×* 150 mm, C18, 3.5 *µ*m [Xbridge (Waters, Milford, MA)]) with a gradient of 115 min length ranging from 97% buffer A (1.25% ammonia in water) to 60% B (1.25% ammonia in 70% acetonitrile in water)^50^.

#### LC-MS/MS analysis

Samples were dried in vacuo and resuspended in 0.05% trifluoroacetic acid in water. Half of the sample was injected into an Ultimate3000 nanoRLSC (Dionex, Sunnyvale, CA) coupled to a Q Exactive or Orbitrap Fusion Lumos (Thermo Fisher Scientific). Peptides were separated on custom-made 50 cm *×* 100 *µ*m (ID) reversed-phase columns (Reprosil) at 55 °C. Gradient elution was performed from 2% acetonitrile to 30% acetonitrile in 0.1% formic acid and 3.5% DMSO over 1 or 2 hours. Q-Exactive plus mass spectrometers were operated with a data-dependent top 10 method. MS spectra were acquired by using 70.000 resolution and an ion target of 3E6. Higher energy collisional dissociation (HCD) scans were performed with 33% NCE at 35.000 resolution (at m/z 200), and the ion target settings was set to 2E5 so as to avoid coalescence^49^. The instruments were operated with Tune 2.2 or 2.3 and Xcalibur 2.7 or 3.0.63.

Synchronous precursor selection (SPS) MS3 experiments were performed on an Orbitrap Fusion Lumos operated with fixed cycle time of 2 sec. MS spectra were acquired at 60.000 resolution and an ion target of 4E5. Tandem mass spectra were acquired in the linear ion trap following HCD fragmentation performed at 38% NCE and an ion target 1E4. MS3 were performed synchronously on the 5 most abundant MS2 signals in the range of 400-2000 m/z using HCD at 70 % NCE in the Orbitrap at a resolution of 30.000 and an ion target of 1E5. The instrument was operated with Tune 2.1.1565.23 and Xcalibur 4.0.27.10. Targeted data acquisition in combination with synchronous precursor selection was performed according to Savitski et al. (2010)^51^, employing the settings described above.

#### Peptide and protein identification

Mascot 2.4 (Matrix Science, Boston, MA) was used for protein identification by using a 10 parts per million mass tolerance for peptide precursors and 20 mD (HCD) mass tolerance for fragment ions. The search database consisted of a UniProt protein sequence database combined with a decoy version of this database created by using scripts supplied by MatrixScience. Carbamidomethylation of cysteine residues and TMT modification of lysine residues were set as fixed modifications. Methionine oxidation, and N-terminal acetylation of proteins and TMT modification of peptide N-termini were set as variable modifications. We accepted protein identifications as follows: (i) For single-spectrum to sequence assignments, we required this assignment to be the best match and a minimum Mascot score of 31 and a 10× difference of this assignment over the next best assignment. Based on these criteria, the decoy search results indicated <1% false discovery rate (FDR). (ii) For multiple spectrum to sequence assignments and using the same parameters, the decoy search results indicated <0.1% FDR. Quantified proteins were required to contain at least 2 unique peptide matches. FDR for quantified proteins was < 0.1%.

#### Isobaric mass tag based quantification

Reporter ion intensities were read from raw data and multiplied with ion accumulation times (the unit is milliseconds) so as to yield a measure proportional to the number of ions; this measure is referred to as ion area^52^. Spectra matching to peptides were filtered according to the following criteria: mascot ion score > 15, signal-to-background of the precursor ion > 4, and signal-to-interference > 0.5 (Savitski et al., 2013) for MS2 based quantification. Fold changes were corrected for isotope purity as described and adjusted for interference caused by co-eluting nearby isobaric peaks as estimated by the signal-to-interference measure^52^. MS3 based quantification was performed identically but excluding the adjustment for interference caused by co-eluting nearby isobaric peaks. Protein quantification was derived from individual spectra matching to distinct peptides by using a sum-based bootstrap algorithm; 95% confidence intervals were calculated for all protein fold changes that were quantified with more than three spectra^52^ for MS2 based quantification. An automatic outlier removal procedure was performed on proteins with at least 5 quantified PSMs (peptide-spectrum matches). Briefly, each PSM and respective quantification values were sequentially excluded from the fold change calculation; in cases where the confidence interval was then improved by at least 30%, the given quantification values were permanently excluded from further analyses.

#### Data Analysis

Raw mass spectra were analyzed by Mascot and IsobarQuant^53^. Downstream analyses were performed with R and Bioconductor^54^. TMT reporter intensities were used to calculate the residual KO expression, i.e. KO line over HAP1 WT intensity ratio, per spectrum. Missing reporter intensities were imputed with zeros. Residual protein levels of exons were calculated by calculating the median of all spectra uniquely mapping to every exon of the protein of interest. Exon annotation was obtained from ENSEMBL using biomaRt^46^ and translated to the corresponding amino acid numbers. The position of the spectra was determined as the middle of the identified peptide. Protein domain annotation was obtained from ENSEMBL using biomaRt^46^. All plots were generated using ggplot2^55^.

### BRD4 immunoprecipitation

BRD4 immunoprecipitations were performed using two antibodies. Antibody 1: Rabbit anti-BRD4 amino acids 1312-1362 (Bethyl Laboratories, #A301985A). Antibody 2: Rabbit anti-BRD4 (Abcam #ab128874). 100 *µ*l AminoLink resin (Thermo Fisher Scientific) was functionalized with antibody. 100 *µ*l HAP1 WT or HAP1 BRD4 KO lysate were added and incubated for 2h at 4 °C. Beads were subsequently washed and eluted by adding SDS sample buffer. BRD4 products were detected by Western blotting using the Rabbit anti-BRD4 (Santa Cruz #sc-48772) antibody.

### Inhibitor and histone tail pulldowns

Inhibitor and histone tail affinity enrichment experiments were performed as described previously^32^. 5 *µ*l NeutrAvidin beads (Thermo Scientific) were functionalized with either 0.02 mM panBET inhibitor, GSK923121, or 0.01 mM H4(1-21)-K5_ac_,K8_ac_,K12_ac_. 100 *µ*l HAP1 WT or HAP1 BRD4 KO lysate were added and incubated for 2h at 4 °C. Competition was performed by adding a final concentration of 10 *µ*M IBET151 or DMSO as control. Beads were subsequently washed and proteins eluted by adding SDS sample buffer. Western blotting was performed using the Rabbit anti-BRD4 (Santa Cruz #sc-48772) antibody.

### Methylation analysis

A wild-type HAP1, DNMT1^5bp^ and DNMT1^20bp^ KO clone were seeded at 0.2 106 cells/well in triplicates (6 wells for DNMT1^20bp^ clone). The next day, cells were treated with 0.1% DMSO except 3 wells for DNMT1^20bp^ clone that were treated with the DNMT1 inhibitor RG-108 (GW546419X) at 5 uM (0.1% DMSO final) for 24h. The following day, cells were harvested and genomic DNA was isolated using the “Blood & Cell Culture DNA Mini Kit” from Qiagen and according to the manufacturer’s instructions. Samples were analysed using the Infinium MethylationEPIC BeadChip (Illumina) according to the manufacturer’s instructions. Raw signal intensities obtained from IDAT files were processed with the R/Bioconductor^54^ package minfi^56^. Probes containing single-nucleotide polymorphisms were removed and subsequently beta values were calculated from raw intensities.

### NGLY1 KO generation

K562 cells were transfected with two gRNAs targeting exon 1 and 3 of NGLY1 and Cas9 plasmid (lentiCas9-Blast) by Nucleofection according to the manufacturer’s protocol (Nucleofector, Lonza). LentiGuide-Puro (Addgene plasmid #52963) and lentiCas9-Blast (Addgene plasmid #52962) were gifts from Feng Zhang^57^. Positive transfected cells were selected and enriched with Puromycin (4 *µ*g/*µ*l) and Blasticitidin (3.5 *µ*g/*µ*l) for 15 days. NGLY1 knock out was confirmed in the cell population by western blotting using the anti-NGLY1 antibody from Sigma (HPA036825). Single GFP negative cells were sorted into a 96 well plate and expanded. NGLY1 KO clones were confirmed by Sanger sequencing.

Exon 1 gRNA: CTTGGAGGCCTCCAAAA

Exon 3 gRNA: TCTGCTACTTCTCTCTA

Exon 8 gRNA: GCAAACATGAAGAGGTGATT

### Deglycosylation assay

The deglycosylation-dependent Venus fluorescence assay was performed as described previously^35^. WT and NGLY1 KO clones were infected or transfected with ddVenus reporter and mCherry expression constructs. Cells were treated with DMSO or 20*µ*M zVAD-FMK for 6h at 37 °C. Subsequently, mean fluorescence of Venus and mCherry were measured by flow cytometry on a BD LSR Fortessa using the HTS attachment. mCherry levels were used to control for transfection efficiency and cell size.

## Data Availability

The transcriptomics and proteomics data of the 19 KO lines shown in Figure 1b and the NGLY1 KO lines were deposited to publicly available repositories. The mass spectrometry proteomics data were deposited to the ProteomeXchange Consortium via the PRIDE^58^ partner repository with the dataset identifier PXD010335. RNA-seq data were deposited in the ArrayExpress database at EMBL-EBI (www.ebi.ac.uk/arrayexpress) under accession number E-MTAB-7061. The data for the remaining 174 KO lines shown in Figure 1a is available upon request.

## Acknowledgements

We would like to thank Dinko Pavlinic, Ferris Jung and Vladimir Benes (EMBL Gene Core Facility) for RNA sequencing, Jürgen Stuhlfauth and team for cell banking, Markus Boesche and team for mass spectrometry analyses Satoko Shimamura for cell culture and sample preparation, and the microarray unit of the DKFZ Genomics and Proteomics Core Facility for providing the Illumina Human Methylation arrays and related services. The NGLY1 work was supported by the Grace Science Foundation.

## Author Contributions

A.H.S. and F.Z. analyzed the data. A.H.S. and D.E. performed transcriptomics experiments. N.Z. performed mass spectrometry measurements. G.J. re-sequenced cell lines and analyzed BRD4 and DNMT1 truncation data. W.F.M., K.T. and H.S. performed all NGLY1 KO experiments and initial analyses. A.M.M. performed the BRD4 functional experiments. P.G., T.B., M.F.S., M.B., L.M.S., W.H. and G.D. supervised the work. A.H.S., F.Z., W.H. and G.D. wrote the manuscript with input from all authors.

