## Supplementary Figures for "Biological Plasticity Rescues Target Activity in CRISPR Knockouts"

Table S1: Chromatograms resulting from Sanger sequencing of the BRD4, DNMT1, and NGLY1 KO lines. Deletion sites are shown by a blue bar.

| Gene | Indel | Sequence and indel position | sgRNA sequence | Primer sequence |
|---|---|---|---|---|
| BRD4 | Δ5 | 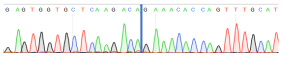 | GAGTGGTGCTCAAGACACTA | TCTGCCAGTAATGGGGGATGG |
| DNMT1$^{20bp}$ | Δ20 | 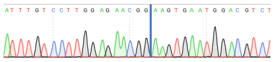 | CGGTGCTCATGCTTACAACC | TCTGACACTTGTTTACATTCACTGC |
| DNMT1$^{5bp}$ | Δ5 and G to A mutation | 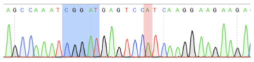 | CTTGATGGACTCATCCGATT | TAGGACTTACAAGATGGCAAGACAA |
| NGLY1$^{c15}$ | Δ26 | 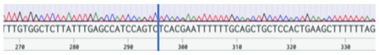 | TCTGCTACTTCTCTCTA | CCCTAATGATGAAAAATATAGATCC |
| NGLY1$^{c20}$ | Δ26 | 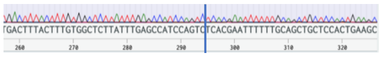 | TCTGCTACTTCTCTCTA | CCCTAATGATGAAAAATATAGATCC |

**a**

| TMT | 126 | 127L | 127H | 128L | 128H | 129L | 129H | 130L | 130H | 131L |
|---|---|---|---|---|---|---|---|---|---|---|
| E.Coli | 0 | 1 | 1 | 0.5 | 0.5 | 0.2 | 0.2 | 0.1 | 0.1 | 1 |
| human | 1 | 1 | 0 | 1 | 0 | 1 | 0 | 1 | 0 | 1 |

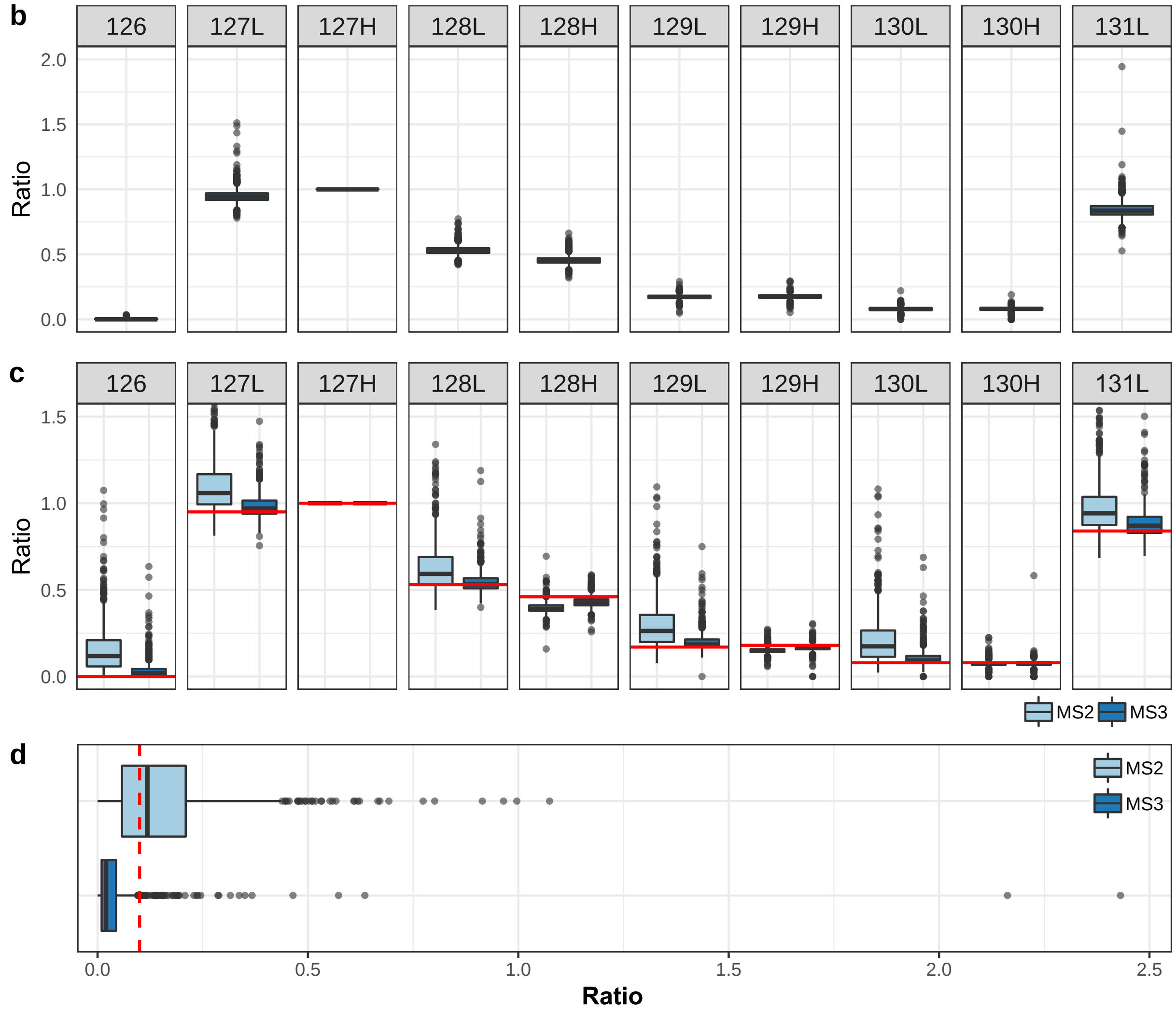


**e**

| Analysis | Proteins | Proteins above 0.1 | FPR |
|---|---|---|---|
| MS2 | 772 | 442 | 0.57 |
| MS3 | 587 | 49 | 0.08 |

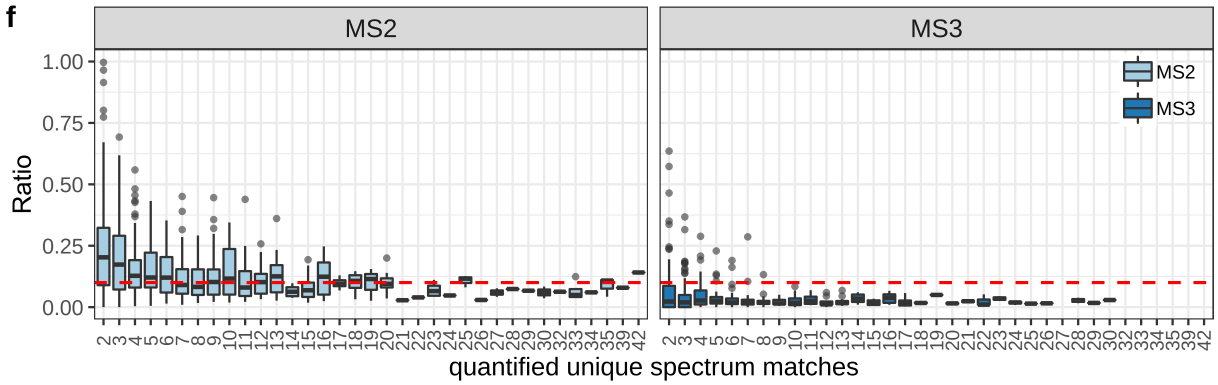


Figure S1: Benchmark experiments to assess ratio compression[52] in MS2 and MS3 analysis with bootstrap based TMT peptide quantification. (a) Design of experiments with and without interference by the complex background. TMT reporter ions were associated to *E. Coli* and human proteins. The table shows the relative amounts of *E. Coli* and human protein in the samples. (Caption continued on next page)

(b) Measured ratios of the respective label versus 127H (the *E. Coli* only control sample) of *E. Coli* proteins analysed without human protein background using MS3 based quantification. (c) Measured ratios with human background, using either MS2 or MS3 analysis. Red bars indicate the expected ratio based on the quantification shown in Panel b. (d) Measured ratios of *E. Coli* proteins in TMT channel 126, which contains only the human background, using either MS2 or MS3 analysis. The red dashed line indicates the value 0.1, which is the cutoff we used for calling residual protein expression in the false positive rate (FPR) calculation in Table S2. (e) Global false positive rate for detection of *E. Coli* proteins, when a threshold of 0.1 is used, as determined from the data for TMT channel 126 using MS2 or MS3 analysis. (f) Observed ratios for TMT channel 126 versus quantified unique spectrum sequence matches per protein using MS2 or MS3 analysis. The red dashed line indicates the value 0.1.

Table S2: False positive rate in individual quantified spectrum sequence match (qusm) bins analysed at a relative fold change of 0.1 for an empty TMT channel (TMT126) of *E. Coli* proteins interfered by the human background analysed by either an MS2 or an MS3 analysis with bootstrap based TMT quantification.

| | **MS2** | | | **MS3** | | |
|---|---|---|---|---|---|---|
| qusm | Proteins | Proteins above 0.1 | FPR in qusm bin | Proteins | Proteins above 0.1 | FPR in qusm bin |
| 2 | 132 | 96 | 0.73 | 113 | 23 | 0.2 |
| 3 | 109 | 74 | 0.68 | 105 | 12 | 0.11 |
| 4 | 101 | 66 | 0.65 | 79 | 5 | 0.06 |
| 5 | 70 | 44 | 0.63 | 43 | 4 | 0.09 |
| 6 | 49 | 27 | 0.55 | 42 | 2 | 0.05 |
| 7 | 57 | 25 | 0.44 | 40 | 2 | 0.05 |
| 8 | 39 | 16 | 0.41 | 33 | 1 | 0.03 |
| 9 | 36 | 19 | 0.53 | 15 | 0 | 0 |
| 10 | 20 | 11 | 0.55 | 20 | 0 | 0 |
| 11 | 18 | 7 | 0.39 | 13 | 0 | 0 |
| 12 | 23 | 12 | 0.52 | 11 | 0 | 0 |
| 13 | 19 | 12 | 0.63 | 14 | 0 | 0 |
| 14 | 10 | 0 | 0 | 8 | 0 | 0 |
| 15 | 16 | 4 | 0.25 | 9 | 0 | 0 |
| 16 | 13 | 7 | 0.54 | 8 | 0 | 0 |
| 17 | 9 | 4 | 0.44 | 10 | 0 | 0 |
| 18 | 7 | 4 | 0.57 | 3 | 0 | 0 |
| 19 | 7 | 4 | 0.57 | 1 | 0 | 0 |
| 20 | 8 | 3 | 0.38 | 2 | 0 | 0 |
| 21 | 1 | 0 | 0 | 2 | 0 | 0 |
| 22 | 1 | 0 | 0 | 3 | 0 | 0 |
| 23 | 5 | 1 | 0.2 | 2 | 0 | 0 |
| 24 | 1 | 0 | 0 | 4 | 0 | 0 |
| 25 | 3 | 2 | 0.67 | 1 | 0 | 0 |
| 26 | 1 | 0 | 0 | 2 | 0 | 0 |
| 27 | 2 | 0 | 0 | – | – | – |
| 28 | 1 | 0 | 0 | 2 | 0 | 0 |
| 29 | 1 | 0 | 0 | 1 | 0 | 0 |
| 30 | 2 | 0 | 0 | 1 | 0 | 0 |
| 32 | 1 | 0 | 0 | – | – | – |
| 33 | 4 | 1 | 0.25 | – | – | – |
| 34 | 1 | 0 | 0 | – | – | – |
| 35 | 3 | 2 | 0.67 | – | – | – |
| 39 | 1 | 0 | 0 | – | – | – |
| 42 | 1 | 1 | 1 | – | – | – |

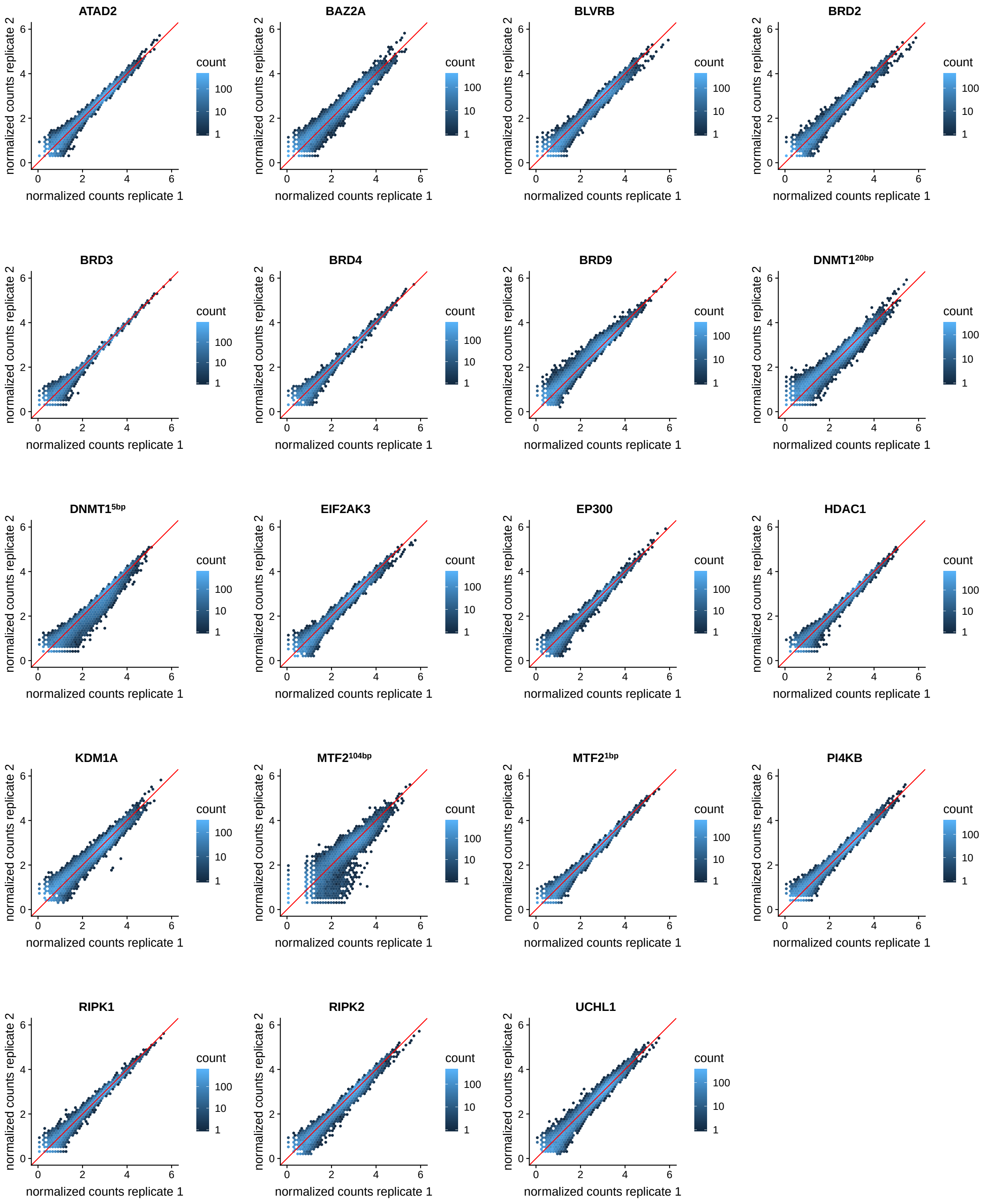


Figure S2: Scatterplots of replicate quantifications from the whole-transcript RNA-sequencing data (see also Fig. 1b). For each KO line, the VST-transformed read counts[44] between two replicates is shown; where more than 2 replicates were acquired, this plot shows the pair with the lowest correlation. The scatterplots use hexagonal binning, with color representing the number of genes falling in each hexagonal bin and the identity line is plotted in red.

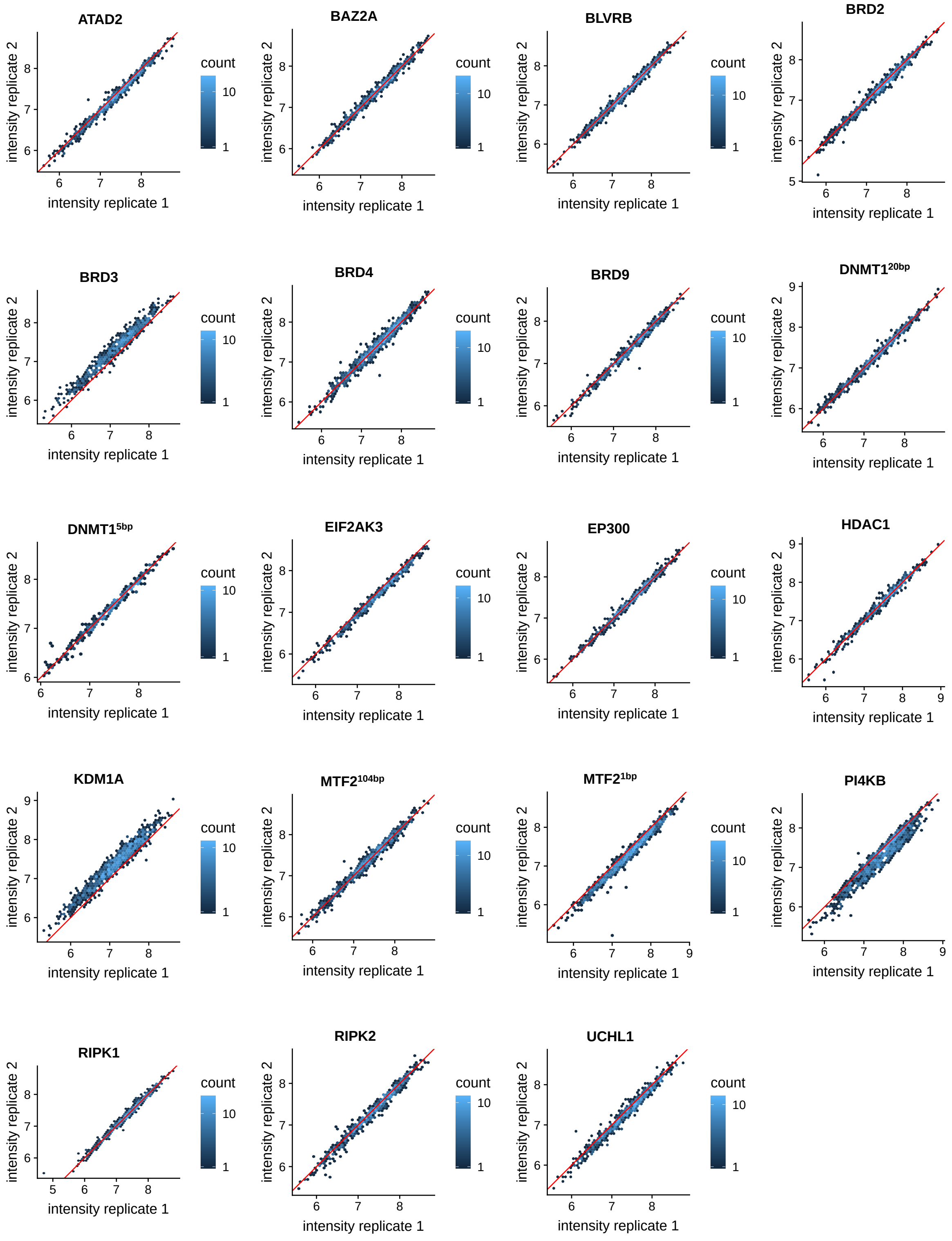


Figure S3: Correlation of replicates for the MS3 proteomics data shown in Fig. 1b. For each KO line, the $log_{10}$-transformed TMT-experiment corrected sumionareas for the pair of replicates with the lowest correlation is shown. The scatterplots use hexagonal binning, with color representing the number of proteins falling in each hexagonal bin and the identity line is plotted in red.
